# Identification and characterization of the HERV-K (HML-8) group of human endogenous retroviruses in the genome

**DOI:** 10.1101/2022.02.10.479833

**Authors:** Mengying Liu, Lei Jia, Hanping Li, Yongjian Liu, Jingwan Han, Xiuli Zhai, Xiaolin Wang, Tianyi Li, Jingyun Li, Bohan Zhang, Changyuan Yu, Lin Li

**Affiliations:** College of Life Science and Technology, Beijing University of Chemical Technology, Beijing 100029, China; Department of Virology, Beijing Institute of Microbiology and Epidemiology, Beijing 100071, China; State Key Laboratory of Pathogen and Biosecurity, Beijing 100071, China

**Keywords:** Human endogenous retrovirus, HML-8, BLAT, GRCh38/hg38, gene regulation

## Abstract

Human endogenous retroviruses (HERV) could vertically transmit in a Mendelian fashion and stable maintenance in the human genome which are estimated to comprise about 8%. HERVs affect human physiology and pathology based on the effect of the provirus-encoded protein or LTR elements. The characterization of the genomic distribution is an essential step to understanding the relationship between endogenous retrovirus expression and diseases. However, the poorly characterization of HML-8 hinders a detailed understanding of the expression regulation of this family in human health and its actual impact on host genomes. In the light of this, the definition of a precise and updated HERV-K HML-8 genomic map is urgently needed. Here we report a comprehensive analysis of HERV-K HML-8 sequences presence and distribution within the human genome, with a detailed description of the different structural and phylogenetic aspects characterizing the group. A total of 40 proviruses and 5 solo LTR elements were characterized with a detailed description of provirus structure, integration time, potentially regulated genes, transcription factor binding sites, and primer binding site feature. The integration time results showed that the HML-8 elements found in the human genome have been integrated in the primate lineage between 23.5 and 52 million years ago (mya). Overall, the results have finally clarified the composition of HML-8, providing an exhaustive background for subsequent functional studies.

**Highlights:**

➢ A comprehensive characterization of the HERV-K (HML-8) in human genome.
➢ There is an apparent preference of HML-8 into intergenic regions and introns.
➢ There are two distinct clusters for the *env* region of the HML-8 elements.
➢ The average time of HML-8 integration in human is 37.1 mya.

## Introduction

Germ cell infections caused by exogenous retroviruses and incorporation into host DNA occurred millions of years ago, leads to the vertical transmission in a Mendelian fashion and stable maintenance of human endogenous retroviruses (HERVs) in the human genome (Bannert and Kurth 2004; Boller *et al*. 2008). HERVs are estimated to comprise about 8% in human DNA (Li *et al*. 2001; Bannert and Kurth 2004). Two hypotheses have been proposed to explain their persistence in the human genome during evolution: parasitic theory and symbiosis theory. In the parasitic theory, HERVs were neutral and rather difficult to eliminate (Doolittle and Sapienza 1980; Weinberg 1980). In contrast, the symbiotic theory held that they were preserved by positive selection (Temin 1992).

HERVs originate as integrated proviruses. A common set of HERV includes the *gag, pro, pol*, and *env* genes which are flanked by two long terminal repeats (LTRs) that act as promoters (Bannert and Kurth 2006). Gene *gag* encodes the structural protein that forms the core of the virion; *pro* encodes the viral protease; *pol* encodes reverse transcriptase and integrase; and *env* encodes the glycoprotein complex that controls receptor-mediated fusion and entry (Johnson 2019).

Even most *gag, pro, pol*, and *env* remain, they are usually inactive due to the accumulation of substitutions, deletions, and insertions. Most HERVs exist in the form of solo-LTRs, produced by homologous recombination between 5[and 3[LTR. LTR is the most volatile sequence in the retroviral genome, with almost no similarities between LTRs of retroviruses from different genera (Benachenhou *et al*. 2013; Johnson 2019). These LTR elements have been shown to influence gene regulation by providing regulatory elements such as enhancers, promoters, splicing sites, and polyadenylation sites for various host genes (Gogvadze and Buzdin 2009; Xue *et al*. 2020).

The classification of HERVs has been controversial for a long time (Blomberg *et al*. 2009). One system exists based on the tRNA molecule, which is used by retroviruses as a primer in replication. For example, the HERV-K element is considered to use lysine tRNA. These naming methods are currently considered incomplete. Another system exists based on phylogenetic methods of the highly conserved *pol* sequence (Cohen and Larsson 1988a; Jern *et al*. 2005). Phylogenetically, HERVs have been divided into three classes, where Class I consists of Gamma retrovirus-like, Class II of Beta retrovirus-like, and of vaguely Spuma retrovirus-like elements (Medstrand *et al*. 2002; Vargiu *et al*. 2016). Thereinto, the HERV-K series, latest acquired by the human species between three and six million years ago, belongs to the Class II of Beta retrovirus-like supergroup (Sverdlov 2000). The groups were initially named HML-1 to HML-6, followed by the definition of HML-7 to HML-10 (Medstrand and Blomberg 1993; Andersson *et al*. 1996; Medstrand *et al*. 1997; Andersson *et al*. 1999). Until 2011, 2 proviruses belonging to the HML-2 branch were identified as new groups HML-11 (Subramanian *et al*. 2011). Currently, HERV-K is divided into subfamilies from HML-1 through HML-11.

HERVs affect human physiology and pathology mainly in two aspects. One aspect is based on the effect of the provirus-encoded protein on the host. The most typical physiological function of HERVs is that Env is highly expressed and involved in the formation of the placenta. Syncytin-1 and 2 are the Env proteins of HERV-W and HERV-FRD, respectively, which maintain the fusion trophoblast cell layer and their connection with the cytotrophoblast layer (Blond *et al*. 2000; Blaise *et al*. 2003). Several proteins encoded by HERV-K are related to cancer, such as germ cell tumors, teratocarcinoma, ovarian cancer, prostate cancer, melanoma, rheumatoid arthritis, and amyotrophic lateral sclerosis (Mameli *et al*. 2017; Arru *et al*. 2018; Garcia-Montojo *et al*. 2018; Arru *et al*. 2021). Np9 promotes the growth of myeloid and lymphoblastic leukemia cells by activation of Notch1, ERK, and AKT pathways through upregulation of β-catenin (Chen *et al*. 2013). The expression of HERV-K Env protein in breast cancer tissue is significantly higher than that in normal breast tissue, which is related to disease progression and poor prognosis (Blaise *et al*. 2003).

The other one is based on LTR elements. They can act as transcriptional regulatory elements to interfere with the expression of upstream and downstream genes. In fact, there are many examples of HERV LTRs acting as new promoters or transcription factor binding sites for genes (Babaian and Mager 2016; Thompson *et al*. 2016; Bannert *et al*. 2018; Fuentes *et al*. 2018). The LTR of HERV-E has been revealed to be located in the upstream of pancreatic amylase gene in the reverse direction, regulating the expression of the amylase gene, and providing promoter activity (Samuelson *et al*. 1990). HERV-K LTR possesses tissue-specific enhancer activity and can be used as the main promoter of the galactopancreatic amylase gene in the human colon and small intestine (Dunn *et al*. 2003).

The transcriptional activation of HERV LTRs also has harmful effects on the body. An in vitro model of human mammary epithelial cell transformation indicated the 5’ LTR promoter activity in tumorigenic cells, suggesting that the cell environment of cancer cells is a key component for inducing the activity of the LTR promoter (Montesion *et al*. 2018). Additionally, HERV-W LTR down-regulated the expression of the GABBR1 gene in schizophrenia (Hegyi 2013). Two members of the HERV-I family induce AZFa gene microdeletions in azoospermia patients (Kamp *et al*. 2000).

For the HERV-K group, it’s known that characterization of the genomic distribution is an essential step to understanding the relationship between endogenous retrovirus expression and diseases. For HML-8, there is one research showing the gene polymorphism of HERV-K11 and indicating that the polymorphisms may arise from an individual-specific basis (Cakmak Guner *et al*. 2018). Besides, there is currently no other information about the characterization of HML-8, which hinders a detailed understanding of the expression regulation of this family in human health and its actual impact on host genomes. In the light of this, the definition of a precise and updated HERV-K HML-8 genomic map is urgently needed.

## Materials and Methods

### 1. HML-8 identification and localization in the human genome (hg38)

To confirm HML-8 provirus and solo LTR localization in the human genome, we selected the Genome Reference Consortium released Dec. 2013 assembly GRCh38/hg38 as the human background sequence and the assembled MER11A- HERVK11-MER11A as a query to perform the HML-8 identification. A traditional BLAT search (Kent 2002) in the UCSC Genome Browser database (Kent *et al*. 2002) was used. DNA BLAT works by keeping an index of the entire genome in memory. The index consists of all overlapping 11-mers stepping by 5 except for those heavily involved in repeats (http://genome.ucsc.edu/cgi-bin/hgBlat).

### 2. Elements distribution prediction and chromosome mapping

To evaluate whether HML-8 is randomly distributed in human chromosomes, we predicted its expected distribution according to the formula: e = Cl ∗ n / Tl (e represents the expected number of integrations in the chromosome, Cl is the length of the chromosome, n is the total identified number of HML-8 loci in the human genome, and Tl is the sum length of all chromosomes) (Grandi *et al*. 2021). The comparison of the actual number of HML-8 loci with the expected elements in the chromosome were analyzed through a chi-square (χ^2^) test.

### 3. Structural Characterization

All 40 HML-8 provirus elements were characterized in detail referring to the Dfam reference MER11A-HERVK11-MER11A by multiple alignments performed with Mega 7 and the subsequent analysis on BioEdit software platform (Hall ; Storer *et al*. 2021). All deletions were annotated.

### 4. Phylogenetic analyses

To confirm the assignment of the identified HML-8 elements, Maximum likelihood (ML) phylogenetic trees were constructed using Mega 7 (Kumar *et al*. 2016). Out of the 40 identified provirus elements, 3 proviruses sequences longer than 80% of the HML-8 reference were used to construct a near-full-length phylogenetic tree. According to the model selection function of Mega 7, the best-fitting model of nucleotide substitution for full-length provirus was GTR+G+I. For the 4 coding regions including *gag, pro, pol*, and *env*, we screened sequences longer than 90% of the corresponding section of HML-8 reference to construct their subregion phylogenetic trees. The best-fitting models of nucleotide substitution for *gag, pro, pol*, and *env* analysis were HKY+G+I, GTR+G (*pro* and *pol*), and HKY+G, respectively. Tree topologies were searched using the NNI procedure. The confidence of each node in phylogenetic trees was determined using the bootstrap test with 500 bootstrap replicates. The final ML trees were visualized using iTOL (Letunic and Bork 2021).

### 5. Estimation of the integration time of HML-8

To estimate the time of integration, we assumed a substitution rate of 0.2%/nucleotides/million years for the human genome and used this rate to assess the action of divergence on each HML-8 sequence (Lebedev *et al*. 2000). Estimation of the integration time is calculated based on the formula T = D% / 0.2%, which T is the estimated time of integration (in million years) and D% is the percentage of divergent nucleotides. The divergence values were estimated by comparing each HML-8 internal element *gag, pro, pol*, and *env* genes and its generated consensus. The final age of each sequence was expressed.

### 6. Function prediction of cis-regulatory regions and enrichment analysis

Non-coding regions typically lack biological functions annotation. To examine the biological meaning of HML-8 solo LTRs, their potential association with the analysis of the nearby genes was performed based on the Genomic Regions Enrichment of Annotations Tool (GREAT) against hg38. (McLean *et al*. 2010). Association rule is as follows: Basal + extension: 5000 bp upstream, 1000 bp downstream, 1000000 bp max extension, curated regulatory domains included.

After screening out regulatory genes by GREAT, we used WEB-based Gene SeT AnaLysis Toolkit (WebGestalt) (Liao *et al*. 2019) to perform functional enrichment analysis which is crucial in interpreting the list of interesting genes. WebGestalt (http://www.webgestalt.org) can use three well-established and complementary methods for enrichment analysis, including Over-Representation Analysis (ORA), Gene Set Enrichment Analysis (GSEA), and Network Topology-based Analysis (NTA). The enrichment method used in the current work is ORA. Parameters for the enrichment analysis: Minimum number of IDs in the category: 5. Maximum number of IDs in the category: 2000. FDR Method: BH. Significance Level: Top 10.

### 7. In silico examination of the conserved transcription factor binding sites

The transcriptional binding sites of HML-8 LTR consensus sequence were predicted on the JASPAR (https://jaspar.genereg.net/) database. The taxon was vertebrates, the species was homo sapiens. We selected ChIP-seq data to predict transcription factors with relative profile score threshold ≥95%. The constructed HML-8 LTR consensus sequence alignment and annotation were performed using Geneious software (Kearse *et al*. 2012).

### 8. Primer binding site feature representation

The composition of the primer binding site (PBS) nucleotide sequence of three full- length proviruses and the HML-8 consensus sequence were analyzed using Mega 7 and BioEdit. The grade of conservation at each position was represented with a logo built from WebLogo at http://weblogo.berkeley.edu (Crooks *et al*. 2004).

## Results

### 1. HML-8 elements identification, localization, and actual distribution in hg38

First, a whole HML-8 element distribution was displayed based on Ensembl (www.ensembl.org) (Figure 1A). In total, we characterized 40 HERV-K HML-8 provirus and 5 solo LTR elements. Each HML-8 element is screened out and named according to the genomic locus of insertion (Table 1-2). It has been identified that the average length of these proviruses is 4875 bp. Among them, 6 sequences are larger than 70% of the full-length HML-8 reference sequence (10485 bp), 16 sequences are between 40-70% in length, the remaining 18 sequences are all less than 40% in length. The length of 5 solo LTRs are about 75% of the representative MER11A. The nucleotide sequence of each element was shown in Supplementary dataset 1.

**Figure 1.**
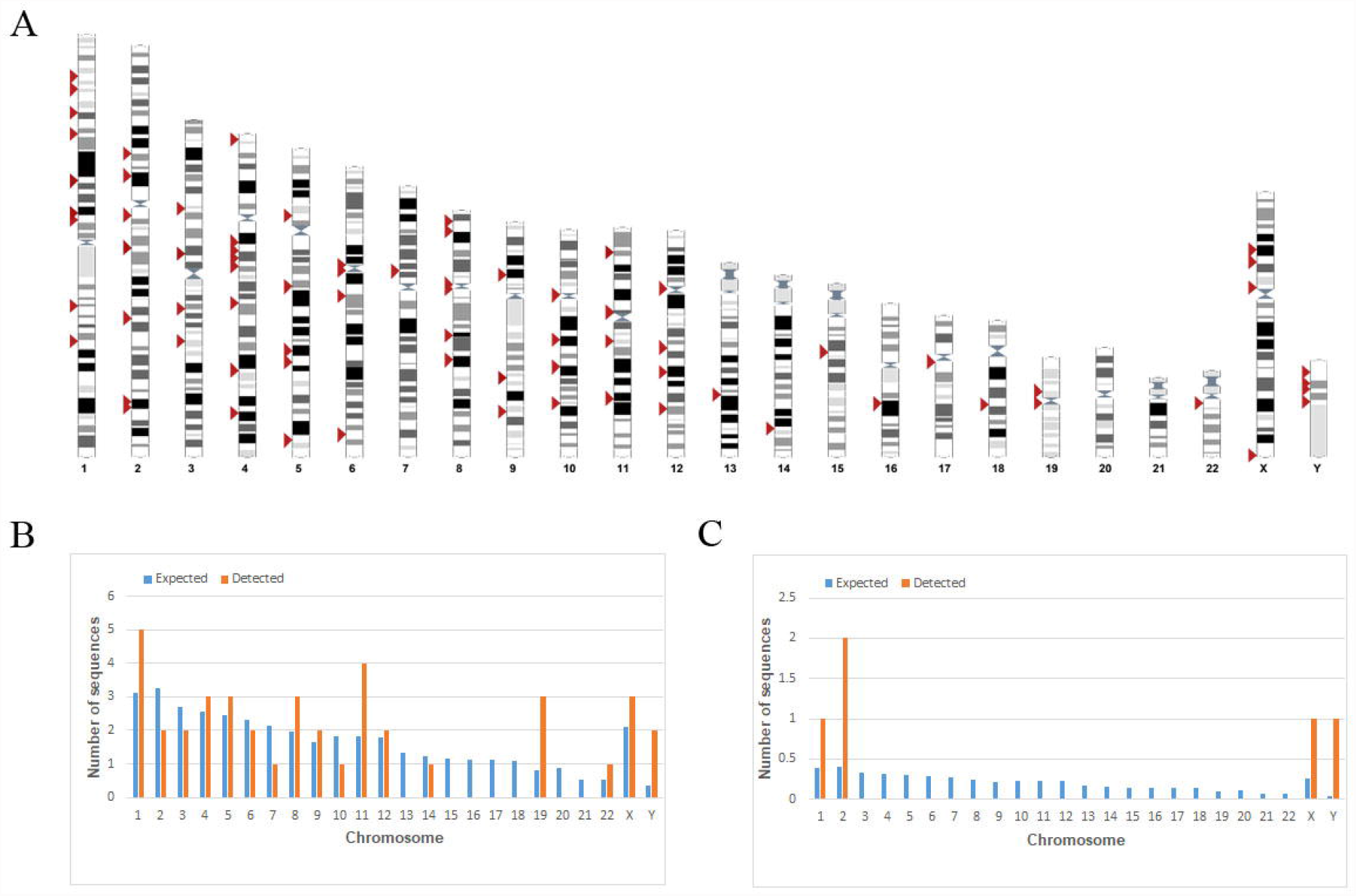
Chromosomal distribution of HML-8 loci. **(A)** All HML-8 elements (red arrows) have been visualized on the human karyotype (www.ensembl.org). The number of HML-8 proviral elements **(A)** and solo LTRs **(B)** integrated into each human chromosome was depicted and compared with the expected number of random insertion events based on chromosomal length. The expected number of sequences in the chromosome were marked with blue and the actual detected number of sequences were marked with yellow.

To assess whether the HML-8 integrations are present in the human genome in a random way, we compare the expected number of integrations with the detected number of HML-8 loci in each chromosome. The results showed that the number of HML-8 integration events observed is always inconsistent with the expected amounts (Figure 1B-C). For the proviral elements, the number of HML-8 insertions in chromosomes 2, 3, 6, 7, 10, 13, 14, 15, 16, 17, 18, 20, and 21 were lower than expected. Thereinto, there were no loci detected in chromosomes 13, 15, 16, 17, 18, 20, and 21 at all. In chromosomes 1, 4, 5, 8, 9, 11, 12, 19, 22, X, and Y, the actual numbers identified are higher than expected (Figure 1B). The solo LTR was only detected in chromosomes 1, 2, X, and Y (Figure 1C). However, the differences were not statistically significant based on the chi-square test. Analysis revealed that HML-8 provirus and solo LTR integration displayed a non-randomly integration way among human chromosomes. Further, all 40 identified provirus elements and 5 solo LTRs were analysed to determine their locations in intergenic regions, intron, or exon (Table 1-2). The results show that 25 provirus elements are located in intergenic regions, accounting for 62.5%. 13 provirus elements are located in introns, accounting for 32.5%. 2 provirus elements are located in both introns and exons, accounting for 5% (Table 1). With respect to solo LTRs, 2 solo LTRs are located in intergenic regions, accounting for 40%. The remaining 3 solo LTRs are located in introns, accounting for 60% (Table 2). The results displayed an apparent insertion preference into intergenic regions and introns.

## Structural Characterization

HML-8 sequences showed a typical proviral structure, with the *gag, pro, pol*, and *env* genes flanked by 5’ LTR and 3’ LTR. According to the annotation information summarized in Dfam database (https://www.dfam.org/family/DF0000189/features), the complete HML-8 contain 4 open reading frames (ORFs). In detail, these structures located in 5’ LTR (from nucleotide 1 to 1266), the *gag* gene (nucleotides 1422 to 3530), the *pro* gene (nucleotides 3341 to 4345), the *pol* gene (nucleotides 4303∼7032), the *env* gene (nucleotides 6890 to 9217), and 3’ LTR (from nucleotide 9220 to10485). To describe the structure of each HML-8 provirus, we aligned 40 HML-8 sequences, annotated the position of the single retroviral component and deletions (Figure 2). In general, there were different degrees of absence of LTR at both ends of all provirus. We obtained 3 relatively complete proviruses (11q22.1, 19p12, 10p11.1), accounting for 80∼90% of complete reference length. However, their LTR structures are still incomplete. More in detail, the integrity of 6 separate regions was summarised in Table 3. It can be noted that many of the proviruses should be retrotransposed pseudogenes because of losing complete LTR structures. Although these genes have been inactivated, they still can provide us with information about the evolutionary history of the gene family or the organism.

**Figure 2.**
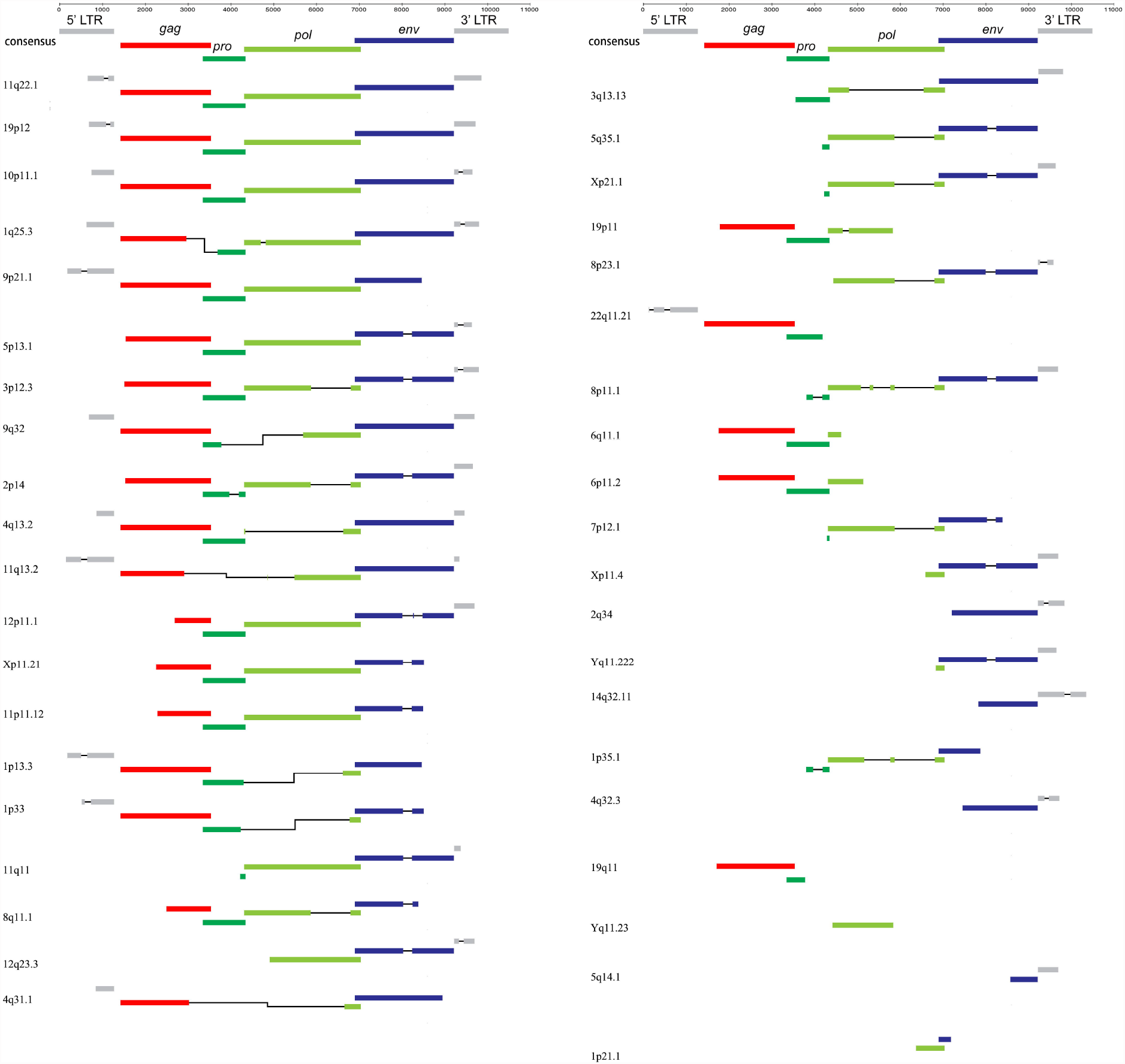
HML-8 proviruses structural characterization. Each HML-8 provirus nucleotide sequence has been compared to the Dfam consensus reference. LTR, *gag, pro, pol*, and *env* regions were annotated. Black lines represented deleted parts.

### 2. Phylogenetic analyses

To confirm the belonging of the newly identified sequence and characterization of the phylogenetic relationships within the HML-8 group, we analyzed 3 proviruses sequences longer than 80% of the HML-8 reference to generate ML trees, including the reference sequences of all Dfam HERV-K groups (HML-1 to 10) and some representative exogenous Betaretroviruses. The results showed that the 3 proviruses all clusters with the Dfam HML-8 group reference with a 100 bootstrap support (Figure 3A).

**Figure 3.**
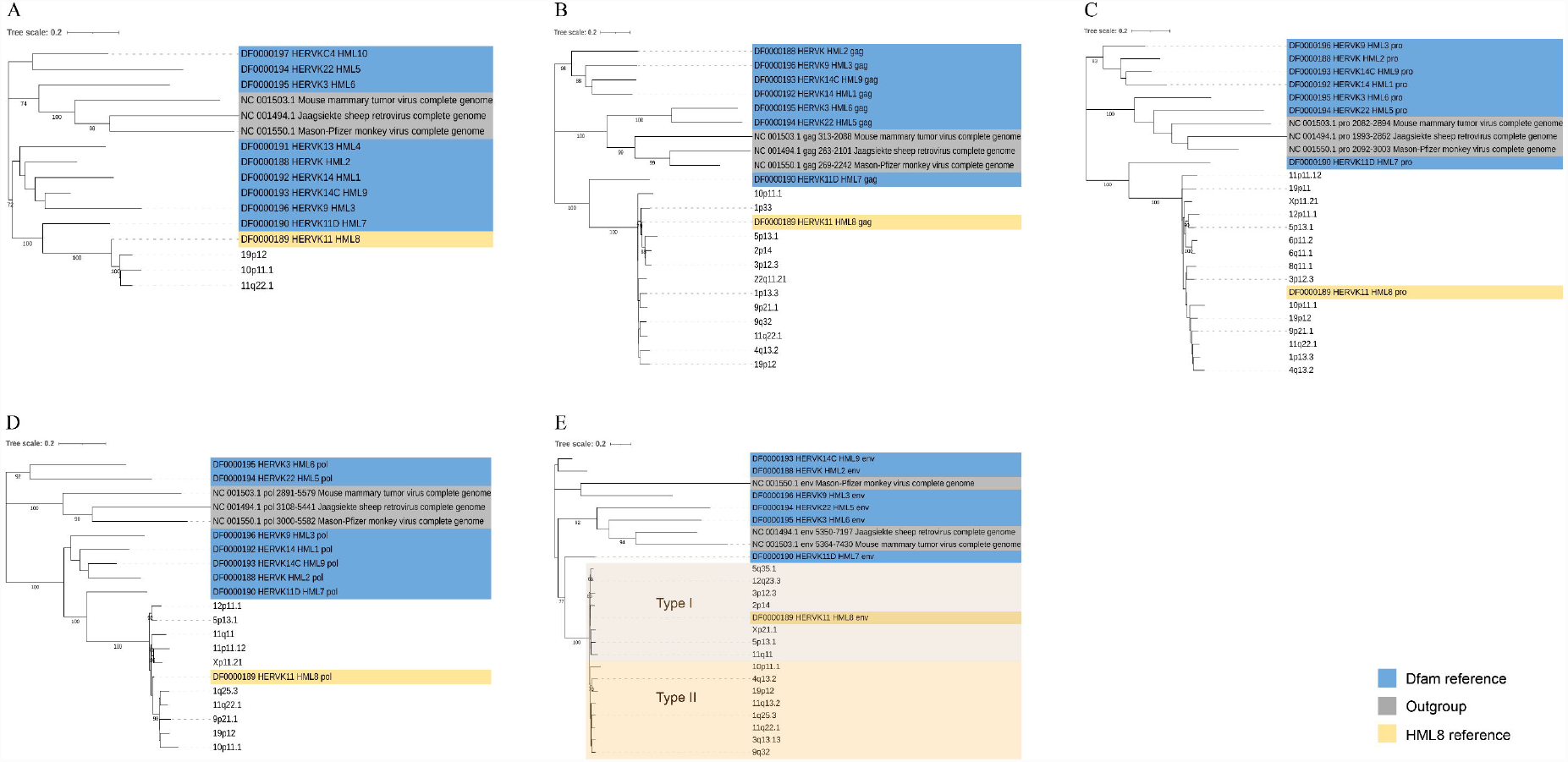
Phylogenetic analysis of the HML-8 near-full-length proviruses and 4 subregions by Maximum Likelihood method. Phylogenetic analyses of HML-8 proviruses **(A)**, *gag* **(B)**, *pro* **(C)**, *pol* **(D)**, and *env* **(E)** together with references. The two intragroup clusters of the *env* gene (I and II) were annotated and depicted with brown and orange background colors, respectively. The resulting phylogeny was tested by using the bootstrap method with 500 replicates. Length of branches indicates the number of substitutions per site.

In addition, we further built the sub-region ML trees for the 12 sequences of *gag* elements, 15 *pro* elements, 10 elements of *pol* elements, and 15 sequences of *env* elements genes, respectively (Figure 3B-E). These phylogenetic groups of different regions of HML-8 all clustered together and were clearly separated from the other HERV-K groups (HML1-7, 9-10). Interestingly, within these main phylogenetic groups, we observed two different clusters located in the *env* section. They were statistically supported by bootstrap values (86% and 76%, respectively) and were named type I and type II. Type I sequences included the Dfam HML-8 *env* reference, whereas Type II elements showed a more divergent structure relative to the group references. The solo LTR of HML1-10 cannot be used as a reference sequence because of its large differences from length to base composition, so we did not draw the phylogenetic tree of HML-8 solo LTR.

### 3. Estimated time of integration

Due to the LTRs of provirus obtained were mostly deleted, we estimated the HML-8 proviruses age based on the *gag, pro, pol, env* regions. Each region longer than 90% of the corresponding references section in length was selected to calculate the integration time. The ancestral sequences of 4 regions have been generated via Mega 7, following the majority rule by the multiple alignments of all corresponding elements. The T value has been estimated by the relation T = D%/0.2%, where 0.2% represents the human genome neutral mutation rate expressed in substitutions/nucleotide/million years. For each region of a provirus, the final T-value was calculated. We provided details on the period of proviruses formation in Table 4. Overall, the results showed that the majority of HML-8 elements found in the human genome have been integrated in the primate lineage between 23.5 and 52 million years ago (mya). The average time of integration was 37.1 mya. Additionally, the integration time of HML-8 elements in chimpanzee was also estimated. The results show it ranges from 15 mya to 52.33 mya (the average time of integration was 35.86 mya), which is consistent with that in humans (data not published).

### 4. Function prediction of cis-regulatory regions and enrichment analysis

The GREAT analysis results were shown in Table 5 which described the associations between each solo LTR and its putatively regulated gene(s). There are a total of 8 genes have been predicted. Among these, one LTR is not associated with any genes. And the other 4 LTRs are associated with 2 genes each (Figure 4A). The absolute distances of these 8 genes to the transcription start site (TSS) are between 5-500 kb (Figure 4B-C).

**Figure 4.**
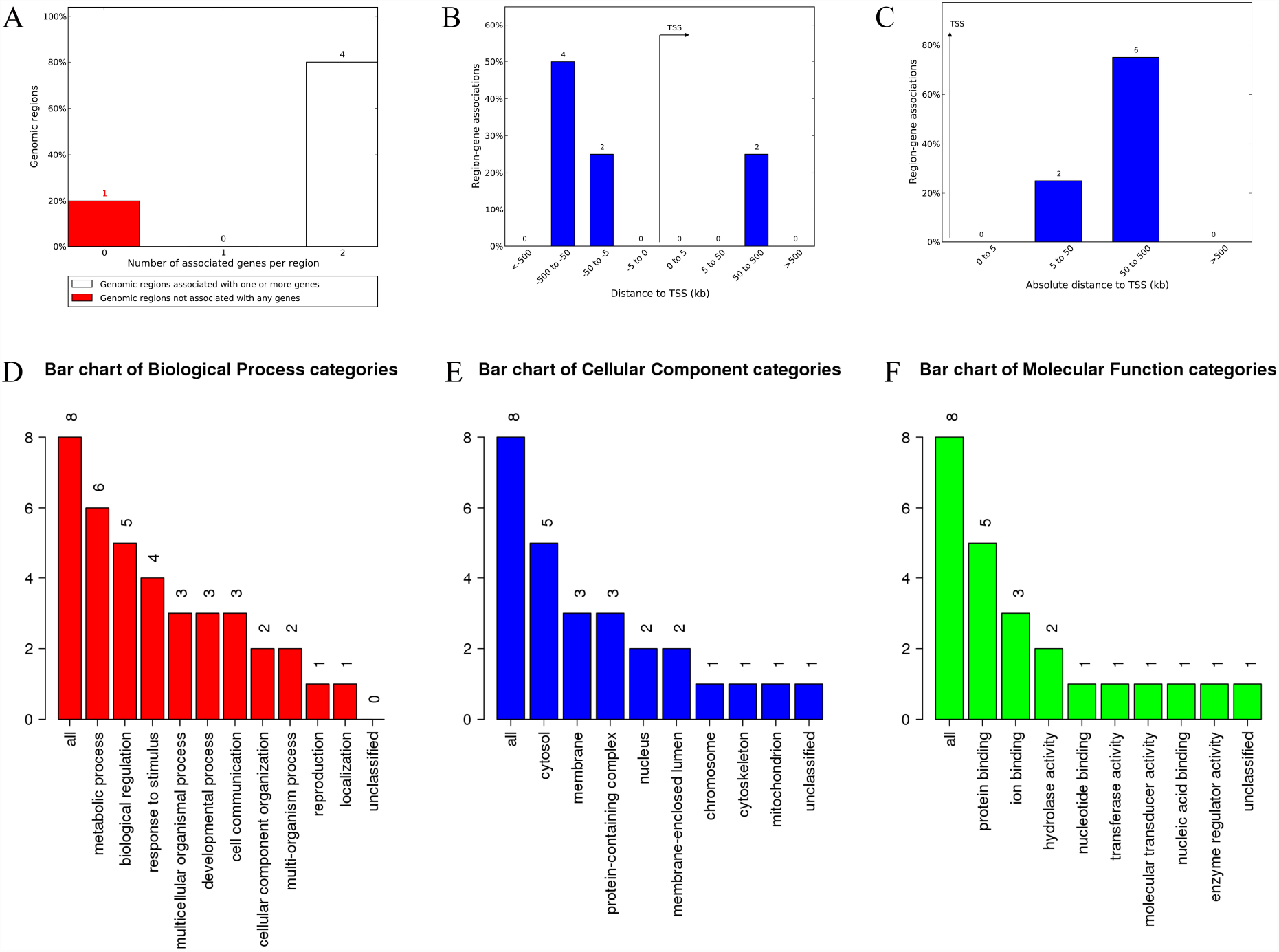
The genes associated with LTRs and GO analysis. **(A)** The number of associated genes per LTR. **(B)** Binned by orientation and distance to TSS. **(C)** Binned by absolute distance to TSS. Each Biological Process **(D)**, Cellular Component **(E)**, and Molecular Function **(F)** category is represented by a red, blue, and green bar, respectively. The height of the bar represents the number of IDs in the gene list and also in the category.

To analyze the biological classification of key genes related to solo LTRs, GO Slim summaries were performed using WebGestalt. Biological processes (BP) analysis revealed that these genes were mainly enriched in the metabolic process and biological regulation (Figure 4D). The changes in cellular component (CC) showed that these genes significantly enriched in the cytosol and the molecular function (MF) enriched in protein binding (Figure 4E-F).

Among 8 unique Entrez gene IDs, 8 IDs are all annotated to the selected functional categories and also in the reference list, which are used for the enrichment analysis. Based on the parameters set in the Method section, 10 categories are identified as enriched categories. As shown in Fig. 5A, a total of 10 enriched categories are identified for Biological Process including carnitine biosynthetic process, positive regulation of DNA ligation, regulation of DNA ligation, amino-acid betaine biosynthetic process, positive regulation of protein deubiquitination, single-strand break repair, regulation of protein deubiquitination, carnitine metabolic process, pentose metabolic process, and negative regulation of fibroblast growth factor receptor signal. The bar chart shows the enrichment ratio of the results. When top results are chosen to be returned and the false discovery rate (FDR) for the categories is ≤0.05, the colors of the bars are in a darker shade than when the FDR exceeds 0.05. The volcano plot in Fig. 5B shows the log2 of the FDR versus the enrichment ratio for all the functional categories in the database, highlighting the degree by which the significant categories stand out from the background. The size and color of the dot are proportional to the number of overlapping (for ORA). The significantly enriched categories are labeled, and the labels are positioned automatically by a force field- based algorithm at startup. The enrichment results for Cellular Component and Molecular Function are illustrated in Supplementary figure 1-2. It must be noted that these results are entirely speculative and that future research is required to confirm any of the implied associations between the solo LTRs and the nearby genes.

**Figure 5.**
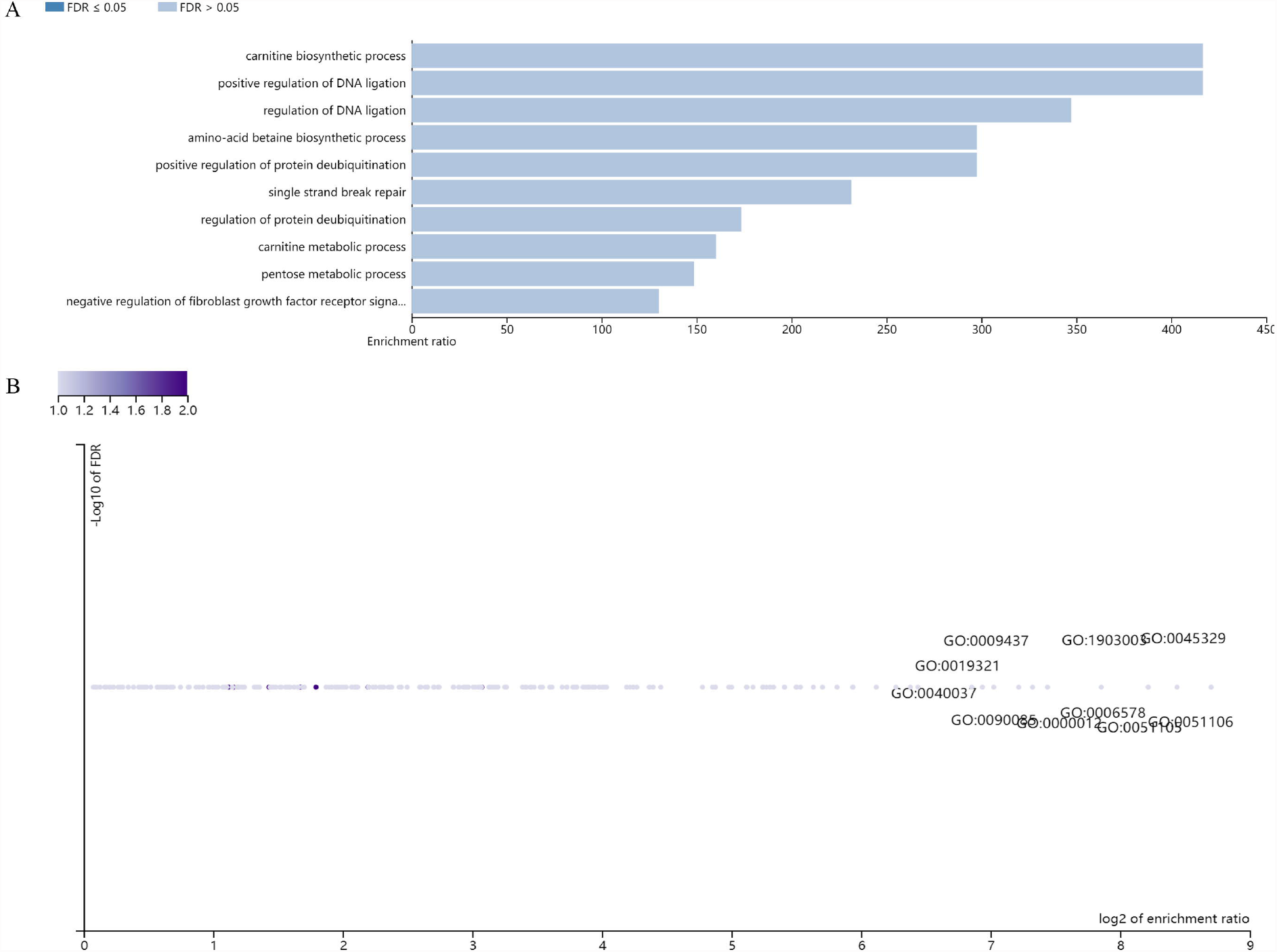
Enrichment result categories binned by Biological Process. **(A)** The bar chart plots the enrichment results vertically with the bar width equal to the enrichment ratio in ORA. **(B)** Customizable volcano plot. Inset shows an initial layout for comparison.

### 5. In silico examination of the conserved transcription factor binding sites

HML-8 exhibiting specific base insertions may influence the complexity of LTR transcriptional regulation (Subramanian *et al*. 2011). A complete view of the putative transcription factor binding sites within the HML-8 LTR were shown in Figure 6A. A total of 33 human transcription binding sites were predicted, including 21 transcription factors: HIF1A, SP1, SP2, SP4, STAT3, GATA2, GATA3, GATA4, MXI1, SOX10, RBPJ, KLF1, KLF5, KLF7, KLF12, PRDM1, CREB1, THAP1, MAZ, ZEB1, THAP1. The motifs are marked on the sense strand and antisense strand.

**Figure 6.**
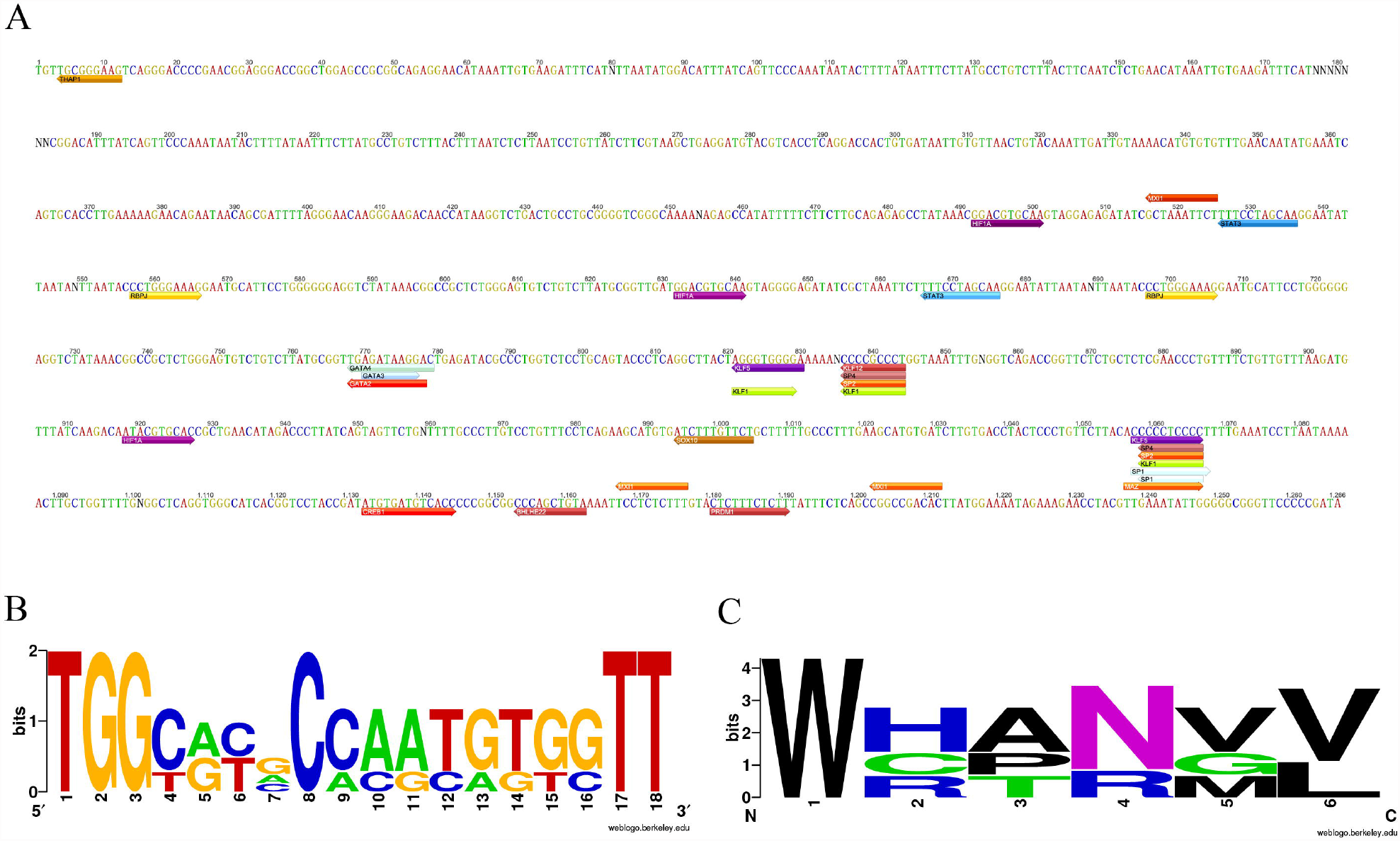
In silico examination of the conserved transcription factor binding sites and logos representing of the PBS of HML-8. **(A)** The forward arrows indicate sense strand, the reverse arrow indicates antisense strand. Different transcription factors are marked with different colors. **(B)** PBS nucleotide sequence; **(C)** PBS amino acid sequence. Created at http://weblogo.berkeley.edu/logo.cgi

### 6. PBS type of HML-8 sequences

Traditionally HERVs have been named per the tRNA that binds their RT enzyme and PBS (Cohen and Larsson 1988b). Thus, HERV-K is named after lysine-tRNA. In the analyzed 3 proviral and consensus sequences of HML-8 elements, the PBS was located approximately from nucleotide 3 to 20 in nucleotides downstream the 5’LTR. To summarize the general variation of the PBS sequence within HML-8 group we generated a logo in which the letter height is proportional to the nucleotides conservation at each position (Figure 6B-C). The result showed the TGG starting nucleotides were the most conserved among the 18 bases analyzed which had a tryptophan (W) PBS type instead of lysine (K), confirming the relatively low taxonomic value of this feature.

## Discussion

Since the discovery of the HERV-K family playing a physiological and pathological role in the body, great attention has been dedicated to further understanding their impact on the host. One of the problems faced in the situation is still the lack of a complete and updated description of the HERV-K sequences in the human genome, their genomic background, and detailed knowledge of HERV-K single members. Until now, the characterization and identification of HML1-6, and HML10 groups have been carried out (Seifarth *et al*. 1998; Mayer and Meese 2002; Lavie *et al*. 2004; Flockerzi *et al*. 2005; Subramanian *et al*. 2011; Grandi *et al*. 2017; Pisano *et al*. 2019). Among HML groups, HML2 is the best-described one, including a full genomic characterization, time of integration, and subtypes (Subramanian *et al*. 2011). The precise identification of these subtypes has led to studies of their roles in physiological tumors or neurological diseases.

In the present study, we completed the identification and characterization of the HML-8 proviruses and solo LTRs in human DNA, providing the first exhaustive description of this group. Following the approach carried in previous published study(Grandi *et al*. 2017; Pisano *et al*. 2019), a total of 40 HERV-K HML-8 provirus and 5 solo LTR elements have been characterized. The chromosomal distribution of these proviruses and the solo LTRs revealed a non-random integration pattern. This bias is probably linked to the strong preference of HML-8 elements to be inserted in proximity or within human genes introns (Vargiu *et al*. 2016). All 40 identified provirus elements and 5 solo LTRs were analysed to determine their locations in intergenic regions, intron, or exon (Table 1-2). The results display an apparent insertion preference into intergenic regions and introns.

Overall, structural characterization revealed that only 3 HML-8 members (7.5%) retained the almost complete proviral structure. The majority of HML-8 elements show a defective proviral structure, with a mean length of 4875 bp and the complete loss of 5’ LTR (70%) and 3’LTR (40%). The deletion of more than 40% of reference length in *gag, pro, pol*, and *env* genes accounted for 52.5%, 50%, 55%, and 30%, respectively.

Phylogenetic analysis showed that 3 sequences of HML-8 near-full-length proviruses as well as 12 sequences of *gag*, 15 elements of *pro*, 10 elements of *pol*, and 15 sequences of *env* form a unique monophyletic cluster, clearly divided from other HML groups and supported by the maximum bootstrap value. The phylogenetic tree of *env* regions revealed the presence of two well-supported clusters, identified here as type I and II and including 7 and 8 members, respectively.

Concerning the time of integration estimation, the traditional approach based on the divergence between the two LTRs of the same provirus was not applicable due to the lack of enough LTR sequences. Therefore, we estimated the HML-8 proviruses age using the *gag, pro, pol*, and *env*. The results indicated the main period of HML-8 integration to be between 23.5 and 52 mya.

A total of 8 genes have been screened out to be potentially regulated by the 5 solo LTRs. GO analyses showed that these genes were mainly enriched in the metabolic process and biological regulation, which indicated these components may be involved in basic biological processes. Through prediction of transcription factors on HML-8 elements by JASPAR, many transcription factors could bind to HML-8 sequence, indicating that it is likely to play a regulatory role in downstream genes. For the analysis of PBS of HML-8, we confirmed the conclusion that this nomenclature is imprecise because HML-8 belong to HERV-K subgroup but its PBS analysis results were tryptophan nevertheless (Blikstad *et al*. 2008). These results are entirely speculative. And experimental validation studies are required to confirm the associations between the solo LTRs and these genes.

## Conclusion

In conclusion, the presented exhaustive characterization of the HML-8 composition and their genomic context of insertion could be the first time to describe this poorly investigated group. These elements could exert in different tissues both in physiological conditions as well as involvement in human disease development. Our study could contribute to better defining their real impact and contribution to our genome.

## List of abbreviations

HERVs: Human endogenous retroviruses
LTR: long terminal repeats
HML: human MMTV-like
ML: Maximum likelihood
GREAT: Genomic Regions Enrichment of Annotations Tool
WebGestalt: WEB-based Gene SeT AnaLysis Toolkit
ORA: Over-Representation Analysis
GSEA: Gene Set Enrichment Analysis
NTA: Network Topology-based Analysis
Mya: million years ago
TSS: transcription start site
BP: biological processes
CC: cellular component
MF: molecular function
FDR: false discovery rate
ORF: Open reading frame

## Declarations

### Ethics approval and consent to participate

Not applicable.

### Consent for publication

Not applicable.

### Availability of data and materials

All data generated or analyzed during this study are included in this published article.

### Author Contributions

Research design: L.L and Y.C. Performed the analysis: L.J, J.H, H.L, X.Z, X.W, Y.L, T.L, Z.B, and J.L. Contributed to the composition of the manuscript: M.L, L.J, and L.L.

## Acknowledgements

This study was supported by the NSFC (31900157).

---

**Supplementary figure 1**. The enrichment results for Cellular Component (A) The bar chart plots the enrichment results vertically with the bar width equal to the enrichment ratio in ORA. (B) Customizable volcano plot. Inset shows an initial layout for comparison.

**Supplementary figure 2**. The enrichment results for Molecular Function (A) The bar chart plots the enrichment results vertically with the bar width equal to the enrichment ratio in ORA. (B) Customizable volcano plot. Inset shows an initial layout for comparison.

**Supplementary dataset 1**. The nucleotide sequence of HML-8 elements

