## Supplementary figures and images for "Identification and characterization of the HERV-K (HML-8) group of human endogenous retroviruses in the genome"

### Supplementary figure 1

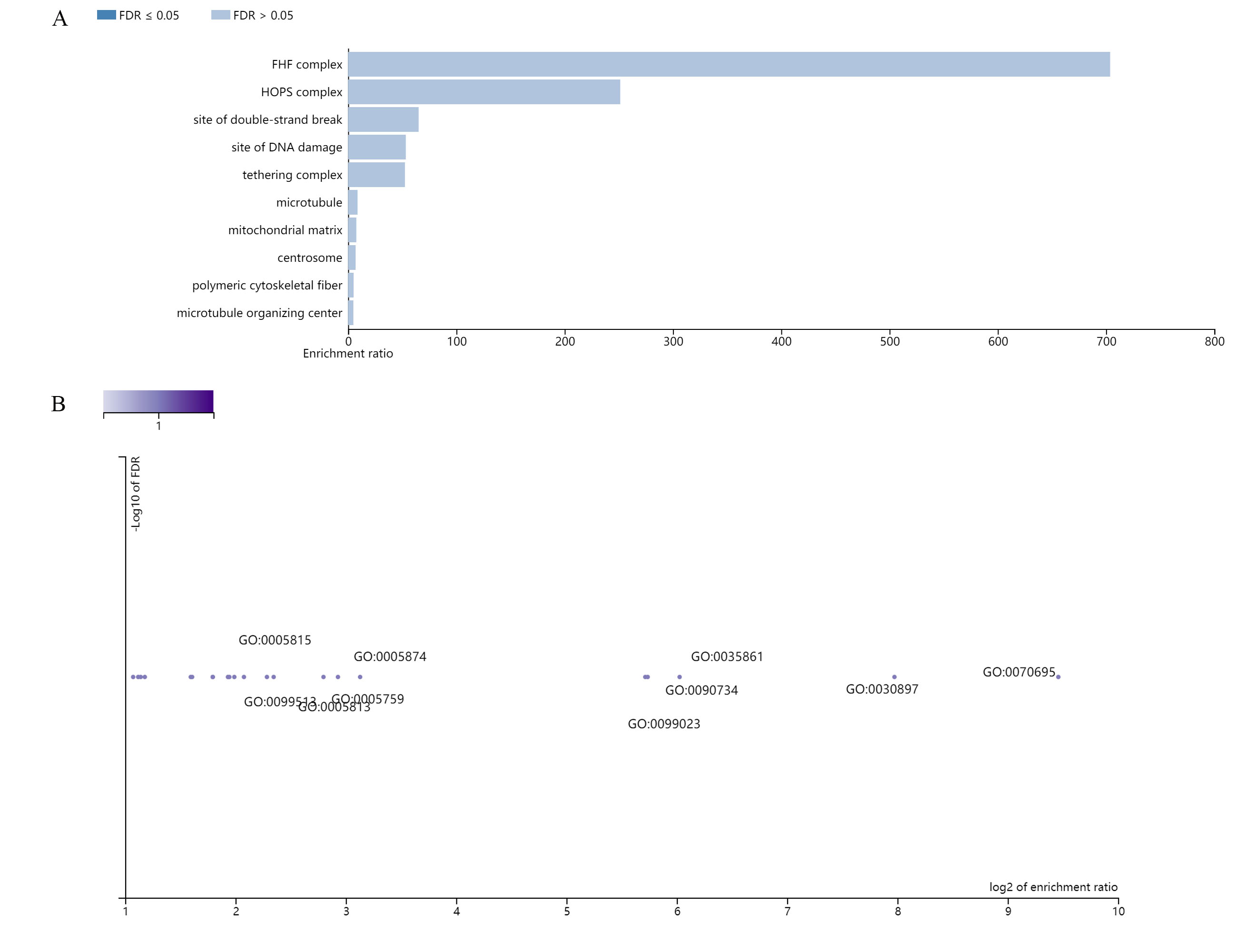

### Supplementary figure 2

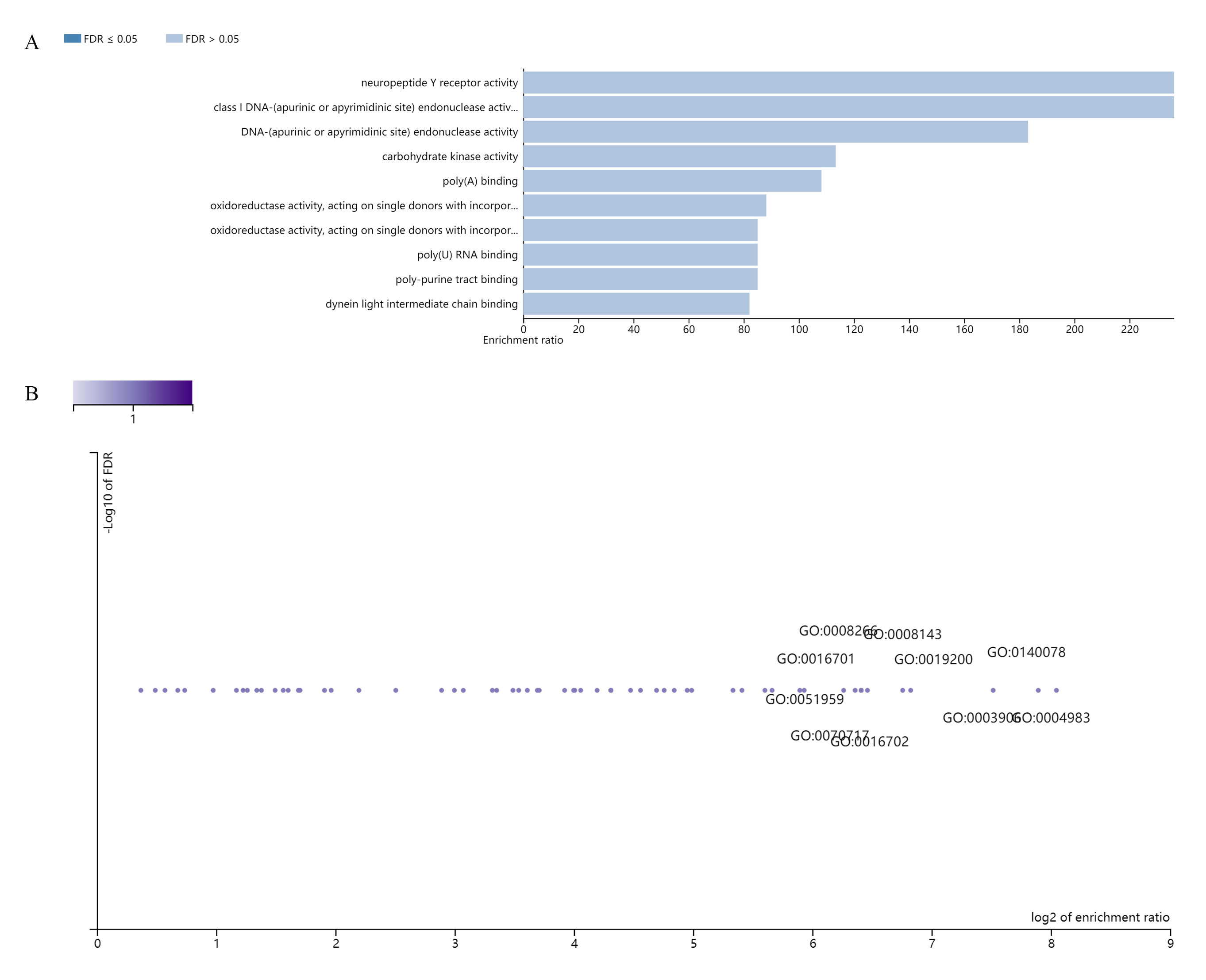
